# Earlier snowmelt increases the strength of the carbon sink in montane meadows unequally across the growing season

**DOI:** 10.1101/2025.04.09.648035

**Authors:** OK Vought, SN Kivlin, HB Shulman, PO Sorensen, DW Inouye, I Ibáñez, P Falb, K Rand, AT Classen

**Affiliations:** Department of Ecology and Evolutionary Biology, University of Michigan, Ann Arbor, MI 48109, USA; Institute for Global Change Biology, School for Environment and Sustainability, University of Michigan, Ann Arbor, Michigan, USA; The Rocky Mountain Biological Laboratory, PO Box 519, Crested Butte, Colorado 81224, USA; Department of Ecology and Evolutionary Biology, University of Tennessee, Knoxville, Tennessee 37996, USA; Natural Resources Science, University of Rhode Island, Kingston, Rhode Island 02881, USA; Climate and Ecosystem Sciences Division, Lawrence Berkeley National Laboratory, Berkeley, California, 94720, USA; Department of Biology, University of Maryland, College Park, Maryland 20742, USA; School for Environment and Sustainability, University of Michigan, Ann Arbor, MI 48109, USA; University of Michigan Biological Station, Pellston, Michigan 49769, USA

## Abstract

1. Warming temperatures are changing winters, leading to earlier snowmelt. This shift can lead to an earlier and potentially longer growing season, which in turn may affect various plant-mediated ecosystem functions. Despite its relevance in the carbon cycle, we still know little about how earlier snowmelt impacts the carbon balance in ecosystems over the growing season, e.g., does it only shift phenology, or does it affect the overall carbon uptake? Most studies rely on interannual variability in snowmelt timing, making it difficult to isolate snowmelt effects from other confounding variables, e.g, temperature and moisture anomalies. To address this uncertainty, we investigated how experimentally advancing snowmelt affects the carbon cycling of montane meadows across the growing season.
2. We experimentally advanced the snowmelt date in a montane meadow by approximately 12 days and collected data every two weeks throughout the growing season, including net ecosystem exchange (NEE), gross primary productivity (GPP), ecosystem respiration (ER), plant composition, and shrub, graminoid, and forb biomass.
3. Early in the growing season, GPP was higher in the early snowmelt plots, though this effect decreased as the growing season progressed. Our modeling of cumulative NEE showed that earlier snowmelt increased the carbon sink strength by 22%, with the strongest effect in the early spring and the effect diminishing as the growing season progressed, with control plots being a greater carbon sink in the later season. Graminoid biomass was 47% higher in plots with earlier snowmelt, but there was no change in total biomass.
4. Synthesis: As winters warm and snowmelt occurs earlier, plant productivity will shift earlier in the growing season, and montane meadows will become a stronger carbon sink. However, this effect will differ seasonally, altering the carbon balance in montane meadows.

## Introduction

In mountain regions, the climate is getting warmer and drier, causing earlier snowmelt (Gottlieb & Mankin, 2024; Lambert et al., 2010; Pepin et al., 2015). Snowmelt timing affects plant phenology and growth in montane meadows (Semenchuk et al., 2016; Vorkauf et al., 2021), as it provides a critical moisture pulse for plant growth in water-limited montane meadows (Ganjurjav et al., 2016; Kawai & Kudo, 2021; Kudo, 2021). Earlier snowmelt can either increase (Zhang et al., 2023) or decrease (Gamon et al., 2013) plant productivity and ecosystem respiration (Li et al., 2016), with a lagged effect on carbon fluxes evident for months (Liu et al., 2023). Earlier snowmelt can also interact with warming temperatures to further dry soils and exacerbate climate change effects on ecosystems (Suzuki, 2014). Changes in plant community composition due to earlier snowmelt can either amplify or mitigate these snowmelt effects on carbon cycling by altering the relative abundance of highly productive species or modifying root production, which influences carbon inputs to the soil through changes in root turnover (D’Imperio et al., 2018; Kelsey et al., 2021; Li et al., 2020; Ma et al., 2024). In systems with large natural snowmelt variation, plant communities are also likely adapted to variable snow conditions, and so they could be less responsive to changing snowmelt (Metz & Tielbörger, 2016). Despite these documented effects, the impact of earlier snowmelt on the carbon balance of montane meadows remains unclear, as most studies rely on natural variations in snowmelt dates rather than experimental manipulations with control plots, which can lead to confounding results, such as temperature and moisture anomalies.

Plants play a crucial role in shaping the response of carbon cycling to changes in snowmelt by altering their phenology, shifting their species composition (Li et al., 2020; Ma et al., 2024), and changing how they allocate resources above and belowground (Darrouzet-Nardi et al., 2019; Sherwood et al., 2017; Vorkauf et al., 2021). With earlier snowmelt, plants often shift their growth earlier in the year, leading to increased productivity in the spring (Blume-Werry et al., 2017; Darrouzet-Nardi et al., 2019; Sherwood et al., 2017; Vorkauf et al., 2021). However, by peak growing season, earlier snowmelt may not significantly change overall biomass (Möhl et al., 2023). Grasses, in particular, tend to respond to changes in snowmelt timing, often becoming more abundant following the pulse of moisture from snowmelt (Ma et al., 2024). Grasses have lower carbon uptake than other plant functional groups (Ibanez et al., 2020), so shifts toward grasses create a positive feedback loop, exacerbating climate change impacts. Whether grasses increase (Li et al., 2020; Ma et al., 2024) or decrease (Liu et al., 2018; Rosbakh et al., 2017) their biomass with less snow and drier conditions depends on the relative rooting depth of co-occurring species and their ability to uptake water. Drier conditions typically favor deeper-rooted species, so species with deeper roots could increase with drought (Liu et al., 2018). Interestingly, changes to snowmelt timing do not typically affect root phenology (Blume-Werry et al., 2017; Darrouzet-Nardi et al., 2019), though experimentally increased snow depth in some ecosystems can increase (Li et al., 2020) or decrease (D’Imperio et al., 2018) root production. These changes in root production can influence carbon uptake, as increased root growth can enhance aboveground carbon uptake, but greater standing root biomass could also release carbon via respiration and increased priming of soil organic matter decomposition (Yan et al., 2023).

Changes in snowmelt timing can affect community and ecosystem-level carbon flux. Earlier snowmelt can increase plant carbon sequestration by enhancing plant productivity earlier in the growing season (Chen et al., 2019; Zhang et al., 2023). However, it may also shorten the growing season if moisture limitations arise later in the growing season (Zona et al., 2022).

While earlier snowmelt could boost gross primary productivity (GPP), it may also increase ecosystem respiration (ER), neutralizing any impact on NEE (López-Blanco et al., 2017). Experimental snow depth studies find mixed effects on carbon uptake, with some studies finding increased uptake (Li et al., 2020) and others finding no effect (Hu et al., 2013). Interestingly, delaying snowmelt had a more significant impact on plant productivity than increasing snow depth, likely because early-season rain diminished the moisture effects of deeper snow (Ma et al., 2024).

To better understand the effects of earlier snowmelt on carbon cycling, we experimentally advanced the snowmelt date in a montane meadow by 12 days. Our results show that shifts in snowmelt timing affect carbon dynamics, as we measured the impacts of earlier snowmelt on plant phenology, biomass accumulation, and carbon fluxes across the growing season in a controlled field experiment. Specifically, we asked: How does an earlier snowmelt date impact the carbon dynamics of a montane meadow across the growing season, considering plant biomass and the overall carbon balance? We hypothesized:

1. Total annual biomass aboveground will not change with snowmelt, as drier conditions later in the growing season would offset any early-season increases. Root production will increase as plants allocate more resources belowground with drier conditions later in the season due to earlier snowmelt.
2. Earlier snowmelt will shift aboveground spring phenology earlier. Root phenology will be unchanged, as root growth may begin before snowmelt.
3. Grasses, which respond to moisture pulses from snowmelt due to their fibrous roots, will be relatively abundant early in the season, especially with earlier snowmelt.
4. NEE sink strength will initially increase with early snowmelt as early-season carbon uptake rises. However, the overall carbon budget will remain unchanged, as early-season productivity gains will be offset by declines in late-season productivity due to soil drying.

## Materials & Methods

### Study sites & experimental establishment

In July 2022, we established a snowmelt manipulation experiment in the East River Meadow near RMBL (38.962396,-106.993848). In a paired design, we established 10, 10 m × 7 m plots to account for spatial differences, with snowmelt manipulation plots paired with control plots (n = 5). The plots all have a similar slope and aspect, with the downslope side facing the East River (west). A minimum of 1.6 m separated each plot. Within each plot, we established two 1 m^2^ subplots in the center zone of the plot, inside a 1 m buffer from the edge of the plot, and marked them to allow repeated measurements in the same location.

The winter of 2022-2023 had a deep snowpack and reached a maximum depth of 246 cm. The total snowfall for the 2022-2023 winter was 921 cm, 12% less than the average from 1974-2022. The majority of annual precipitation is received as snow during the winter (Hubbard et al., 2018). To perform the snowmelt manipulation, we used black shade cloths that covered the entirety of the treatment plots (10 m × 7 m) attached to corner snow poles (Blume-Werry et al., 2017). We deployed the shade cloths on April 20th, 2023, and removed them on May 16th, 2023, when the treatment plots were snow-free. On average, the control plots melted out on May 28th, 2023, indicating that the treatment accelerated snowmelt by approximately 12 days. The snowmelt date has advanced by about 3.5 days per decade at this site since 1975 (Iler et al., 2013). We took biweekly carbon flux and plant measurements from early June to mid-August 2023. The growing season starts when the snow melts in late May and ends when plants senesce in September.

### NDVI, air temperature, & soil moisture measurements

We measured NDVI using Red and Near-IR reflectance measurements from a multispectral camera system (Micasense Altum v1) mounted on an uncrewed aircraft system (UAS, DJI m300 RTK). The instrument was flown on parallel flight lines with 28 m spacing at a height of 100 m above ground level, yielding images with approximately 85% forward overlap and 75% side overlap. We mosaiced, georeferenced, and radiometrically calibrated images from each flight using photogrammetry software (Agisoft Metashape version 1.7) and a standard protocol utilizing ground control points and a reflectance calibration panel. We computed NDVI [(NIR-Red)/(NIR+RED)] at 5 cm resolution for each mosaic before mean-aggregating values to 1 m resolution for time-series analysis. NDVI values for the inner 9 m × 6 m of the plot were then averaged to calculate one NDVI value per plot per UAS flight. Flights were conducted near midday (10 am - 3 pm local time) in sunny conditions approximately once per week during our sampling period.

To measure air and soil temperature (°C) and soil moisture (% volumetric soil water content, VWC), we used TMS-4 sensors (TOMST, Praha, Czech Republic). We installed a single TMS-4 probe in the summer of 2022 in the center of each plot. We collected air temperature at 15 cm above the soil surface and soil temperature at 6 cm below the soil surface level every 15 minutes throughout the growing season. We also measured soil moisture using TMS-4 probes across a 0-15 cm depth every 15 minutes (Wild et al., 2019).

### Plant community & plant biomass measurements above and belowground

We quantified plant community composition and biomass using the pin-drop method (McLaren & Turkington, 2011). We placed a 0.5 m^2^ frame on each subplot and dropped a pin at 25 evenly spaced points, recording the type of plant and the number of times the pin hit to measure plant diversity and to calculate plant biomass. To calculate plant biomass, we built allometric equations that correspond pin-drop hits to aboveground plant biomass. To collect biomass data, we set up 15-20 0.5 m^2^ plots outside the experimental plots. Within each biomass removal plot, we performed a pin-drop measurement as described above. Then, we clipped all the aboveground plant biomass at ground level, separated it to the species level, dried it, and weighed it. We collected plant biomass data bi-weekly, six times across the growing season, corresponding with sampling of carbon fluxes and plant biomass to account for the seasonal changes in plant biomass. We made one equation to estimate forb biomass, one for graminoid biomass, and another for shrub biomass (plant classifications in Supplementary Table S1). To make each equation, we built a linear regression between the number of hits and biomass for all functional groups separately. In the equation, we included all the different time points to account for how biomass would change over the growing season and included time as a random effect in the model. The forb equation was y = 0.93 + 0.38, and the relationship had an *r*^2^ value of 0.66. The shrub equation was y = 2.26 + 4.52, and the relationship had an *r*^2^ value of 0.63. The graminoid equation was y = 0.58 + 0.387, and the relationship had an *r*^2^ value of 0.58. The in the equations represents the number of pin-drop hits, and y is the total aboveground biomass. We estimated the relative abundance of each functional group pre-treatment and found no difference in percent cover of the functional groups between control and earlier snowmelt plots (Figure S1).

To estimate root biomass, we randomly collected a bulk soil core (0-20 cm) using a hammer corer (6 cm in diameter) from each plot (n = 5), positioned at least 1 m inside the plot. We sieved the soil to 2 mm to sort the roots from the soil core. We then spread the sieved rocks, roots, etc., onto a clean surface and picked out the roots for 15 minutes per sample. We dried the roots at 60° C for 48 hours and weighed them to determine root biomass. We sampled nine times throughout the growing season.

### Carbon flux measurements

To measure how earlier snowmelt impacted carbon fluxes, we measured net ecosystem exchange (NEE) and ecosystem respiration (ER) six times throughout the growing season and calculated gross primary productivity (GPP) as the sum of those measurements. We measured NEE in the subplots using an infrared gas analyzer (Li-Cor, Lincoln, Nebraska, USA) inside an airtight, translucent chamber with a volume of 0.6m^3^ and an area of 1m^2^ (Prager et al., 2021). The chamber was sealed to the ground with chains to ensure it was airtight. We took the carbon measurements for 90 seconds at four different light regimes to calculate NEE inside the chamber. We used full sun where the photosynthetic photon flux density (PPFD) was greater than 1000 μmol m s, full darkness where the PPFD was zero μmol m s, and two in-between light levels. Full darkness is a measurement of ER. To block the light, we used one or two layers of plastic shade cloth, each reducing the PPFD by ∼50% inside the chamber (i.e., to about 25% of ambient light with two layers). To measure PPFD, we used an MQ-100 Apogee PAR meter (Apogee Instruments, Logan, UT, USA), which was mounted in the center of the chamber. for each light measurement, we plotted the change in carbon dioxide concentration over time to calculate the rate of change (*dC/dT*) and removed any non-linear portions at the beginning and end of the measurement period. To calculate NEE (*µ*mol·m^-2^·s^-1^) from the flux measurements, we used the following equation (Prager et al., 2021) after we converted the carbon dioxide concentrations to dry molar fractions:

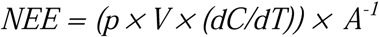

In this equation, *p* is the air density (mol air/m^3^) calculated from *P/RT*, where *P* is the atmospheric pressure (Pa), *R* is the universal gas constant (8.314 J·air K^-1^·mol^-1^), and *T* is the air temperature (K). *V* is the volume of the chamber (m^3^), *dC/dT* is the rate of change of carbon dioxide taken from the chamber (µmol·CO_2_ mol^-1^·air s^-1^), and *A* is the ground area of the chamber (m^2^). We estimated NEE at PPFD = 800 μmol m^-2^ s^-1^ (NEE_800_) to standardize our measurements over time and space and allow us to directly compare carbon dioxide fluxes between the treatments. We estimated NEE_800_ by creating a light response curve with NEE against PPFD and fitting a line between our four light measurements. We then solved the equation for PPFD = 800 μmol m^-2^ s^-1^ (Sundqvist et al., 2020). GPP_800_ is then calculated using the equation:

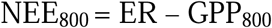

with a negative NEE value indicating net CO_2_ uptake and a positive NEE indicating net CO_2_ release to the atmosphere.

### Data analysis

To determine the effect of earlier snowmelt on total aboveground biomass, forb biomass, shrub biomass, graminoid biomass, root biomass, NEE, GPP, and ER throughout the growing season, we ran linear mixed effect models using the package *nlme* (Pinheiro et al., 2023; Pinheiro & Bates, 2000) in R, version 4.15 (R Core Team, 2023). We ran an identical linear mixed-effects model for all the analyses. We included the interaction between the treatment and date (as day of the year, a continuous variable) as fixed effects. The plot was included as a random effect, and we included a correlation structure to account for temporal autocorrelation. To determine the effect of earlier snowmelt on total aboveground biomass, forb biomass, shrub biomass, graminoid biomass, root biomass, NEE, GPP, and ER at each measurement point throughout the growing season, we calculated the standardized mean difference (SMD) using Hedge’s *d* metric of effect size using the package *effsize* (Torchiano, 2020) in R, version 4.15 (R Core Team, 2023). If SMD > 0, earlier snowmelt increased the variable (a positive effect), if SMD < 0 (a negative effect), earlier snowmelt caused a decrease in the variable. To measure the effect of plant biomass (graminoid, shrub, and forb) on carbon fluxes (NEE, GPP, and ER), we ran linear mixed effect models using the package *nlme* (Pinheiro et al., 2023; Pinheiro & Bates, 2000) in R, version 4.15 (R Core Team, 2023). We included the interactions between treatment, time, and each plant functional group to measure if the relationships changed with earlier snowmelt or at different times across the growing season.

### Carbon analysis

We analyzed how NEE changed each day throughout the growing season (from May 28th - August 19th) in the control and earlier snowmelt plots. We analyzed NEE at any particular time point as a function of the light (PPFD) when NEE was measured, average soil moisture (*Soilm_i_)*, and air temperature (*Airtemp_i_*) in the plot three days before the NEE measurement was taken, and included an intercept. We estimated distinct light, soil moisture, and air temperature, and intercept parameters for the control and earlier snowmelt plots. We estimated different parameter values for light, soil moisture, and air temperature for the early (May 28th - June 24th), peak (June 25th-July 22nd), and late season (July 23rd-August 19th). Season was determined based on NDVI (Figure 1), early season was before peak greenness, mid season was during peak greenness, and late season was after peak greenness. For each time during the season, NEE at plot *i* was estimated using the following normal likelihood:

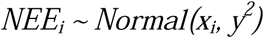

**Figure 1.**
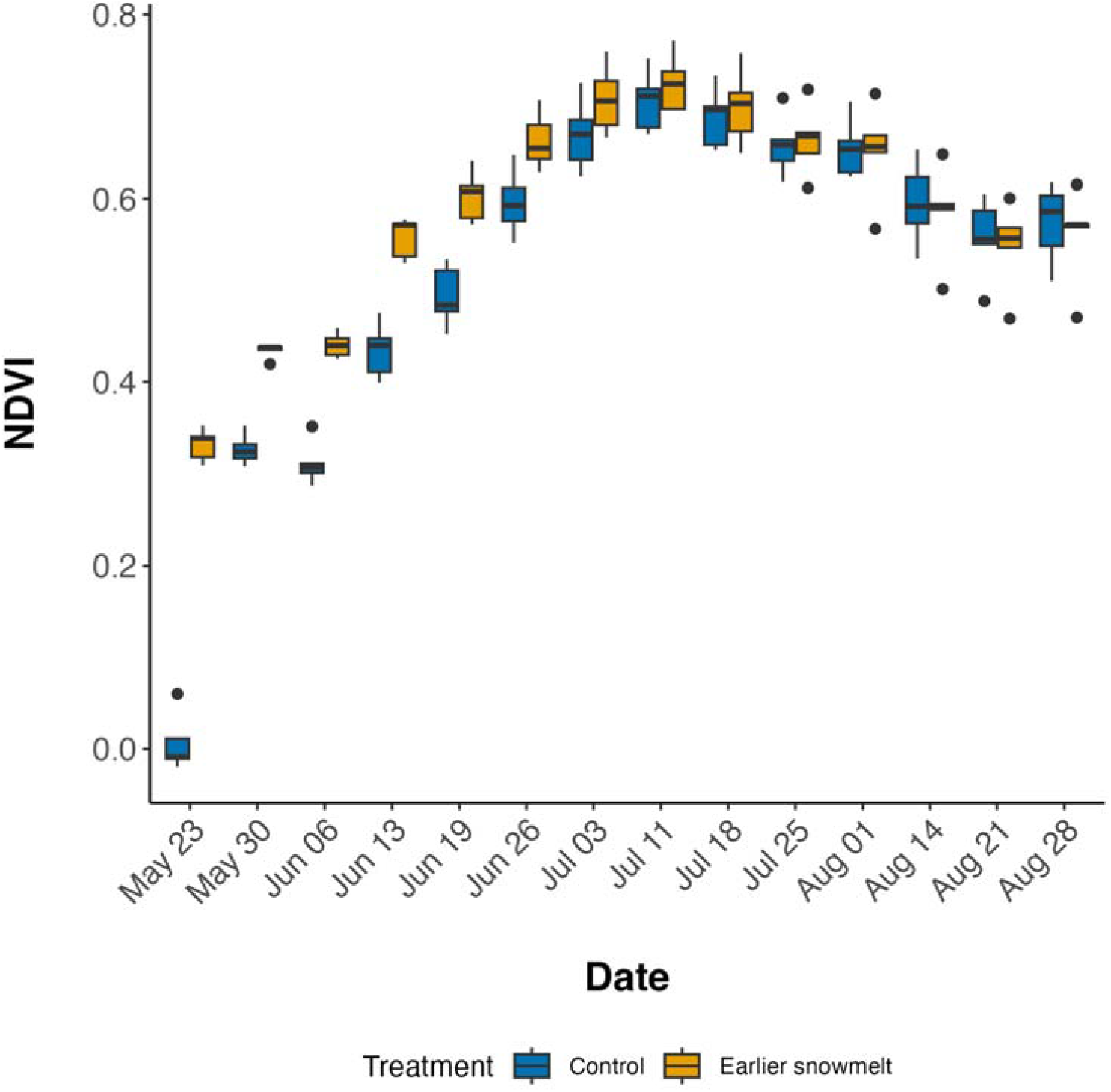
NDVI across the growing season in control and earlier snowmelt plots. For each box, the center line represents the median, and the box represents the interquartile range, with the bottom of the box the 25th percentile, and the top the 75th percentile. The whiskers extend to the largest and smallest data points within 1.5 times the interquartile range.

Process model:

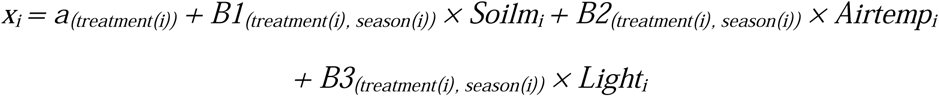

We estimated the parameters using a Bayesian approach from uninformed prior distributions, *a_*_∼ Normal(0, 0.0001), B*_*,*_ ∼ Normal(0, 0.0001),* and *1/y^2^∼ Gamma(0.0001, 0.0001).* We then used the fitted parameter estimates (*a, B1, B2,* and *B3*) and their associated variance to estimate NEE across the growing season as a function of the environmental conditions on each day. The average light (PPFD) per day was estimated for all our plots from the National Solar Radiation Database (Sengupta et al., 2018). NSRD outputs light as Global Horizontal Irradiance, which we converted to PPFD by multiplying by 4.57 (Sager & McFarlane, 1997). Soil moisture and air temperature averages, calculated each day for each plot, and light were used as input data to predict NEE values throughout the growing season. We then summed the daily NEE estimations to calculate total growing season NEE, as well as early, mid, and late season NEE, and included the standard deviation. We used JAGS software (Plummer, 2003) and the *rjags* package in R (Plummer, 2023) to run the analysis. We ran the model for 10,000 iterations following an initial burn-in period of 10,000 iterations. We used three MCMC chains and monitored their mixing to determine convergence. Each model parameter had well-mixed chains and a smooth and even posterior distribution after 10,000 iterations. The goodness of fit was calculated as the *r*^2^ between predicted and observed NEE values to check model fit. All parameter values (means, SDs, and 95% CIs) are in Supplementary Table S2.

## Results

Earlier snowmelt had a minimal effect on soil moisture early in the season but resulted in an 11% decrease in soil moisture from mid-June to mid-August (treatment × time, Figure S2, Table 1).

**Table 1.** Linear mixed effect model results for aboveground biomass, belowground biomass, GPP, ER, and NEE, with a bolded p-value indicating a significant effect.

| Response | Treatment |  |  | Time |  |  | Treatment × Time |  |  |
| --- | --- | --- | --- | --- | --- | --- | --- | --- | --- |
|  | df | F | p | df | F | p | df | F | p |
| Total biomass | 1,18 | 0.0335 | 0.8568 | 1,98 | 36.7935 | **<0.0001** | 1,98 | 2.1718 | 0.01438 |
| Shrub biomass | 1,9 | 1.0411 | 0.3342 | 1,49 | 22.4280 | **<0.0001** | 1,49 | 2.3947 | 0.1282 |
| Forb biomass | 1,18 | 0.0398 | 0.8441 | 1,97 | 14.5089 | **0.0002** | 1,97 | 0.9353 | 0.3359 |
| Graminoid biomass | 1,18 | 1.6902 | 0.2100 | 1,94 | 18.5403 | **<0.0001** | 1,94 | 0.3538 | 0.5534 |
| Root biomass | 1,8 | 0.2260 | 0.6472 | 1,75 | 0.2037 | 0.6531 | 1,75 | 0.5689 | 0.4531 |
| NEE | 1,18 | 0.2036 | 0.6572 | 1,98 | 20.9097 | **<0.0001** | 1,98 | 1.9490 | 0.1658 |
| GPP | 1,18 | 0.0185 | 0.8934 | 1,98 | 17.1742 | **0.0001** | 1,98 | 4.9033 | **0.0291** |
| ER | 1,18 | 0.0869 | 0.7715 | 1,98 | 0.1040 | 0.7478 | 1,98 | 2.2621 | 0.1358 |
| Soil moisture | 1,8 | 0.8500 | 0.3846 | 1,20148 | 149883 | **<0.0001** | 1,20148 | 652.37 | **<0.0001** |
| Air temperature | 1,8 | 0.1510 | 0.7079 | 1,20148 | 395.02 | **<0.0001** | 1,20148 | 0.0930 | 0.7601 |
| NDVI | 1,8 | 9.1886 | **0.0163** | 1,128 | 73.002 | **<0.0001** | 1,183 | 12.229 | **0.0006** |

Earlier snowmelt did not significantly affect air temperature (Figure S2, Table 1). Soil moisture and air temperature both changed significantly over time, reflecting seasonal trends, with conditions becoming drier and warmer as the growing season progressed (Figure S2, Table 1). NDVI was higher in the earlier snowmelt plots early in the season but became more similar to the control plots as the season progressed (treatment × time, Figure 1, Table 1).

Total aboveground biomass increased during the growing season, ranging from an average minimum of 110 g/m^2^ in early June to a maximum of 183 g/m^2^ in mid-July. However, earlier snowmelt did not significantly affect the total aboveground biomass (Figure 2, Table 1). Root biomass varied within each sampling point, between treatments, and throughout the growing season, with earlier snowmelt plots showing an advancement in peak root biomass (Figure 2, Table 1).

**Figure 2.**
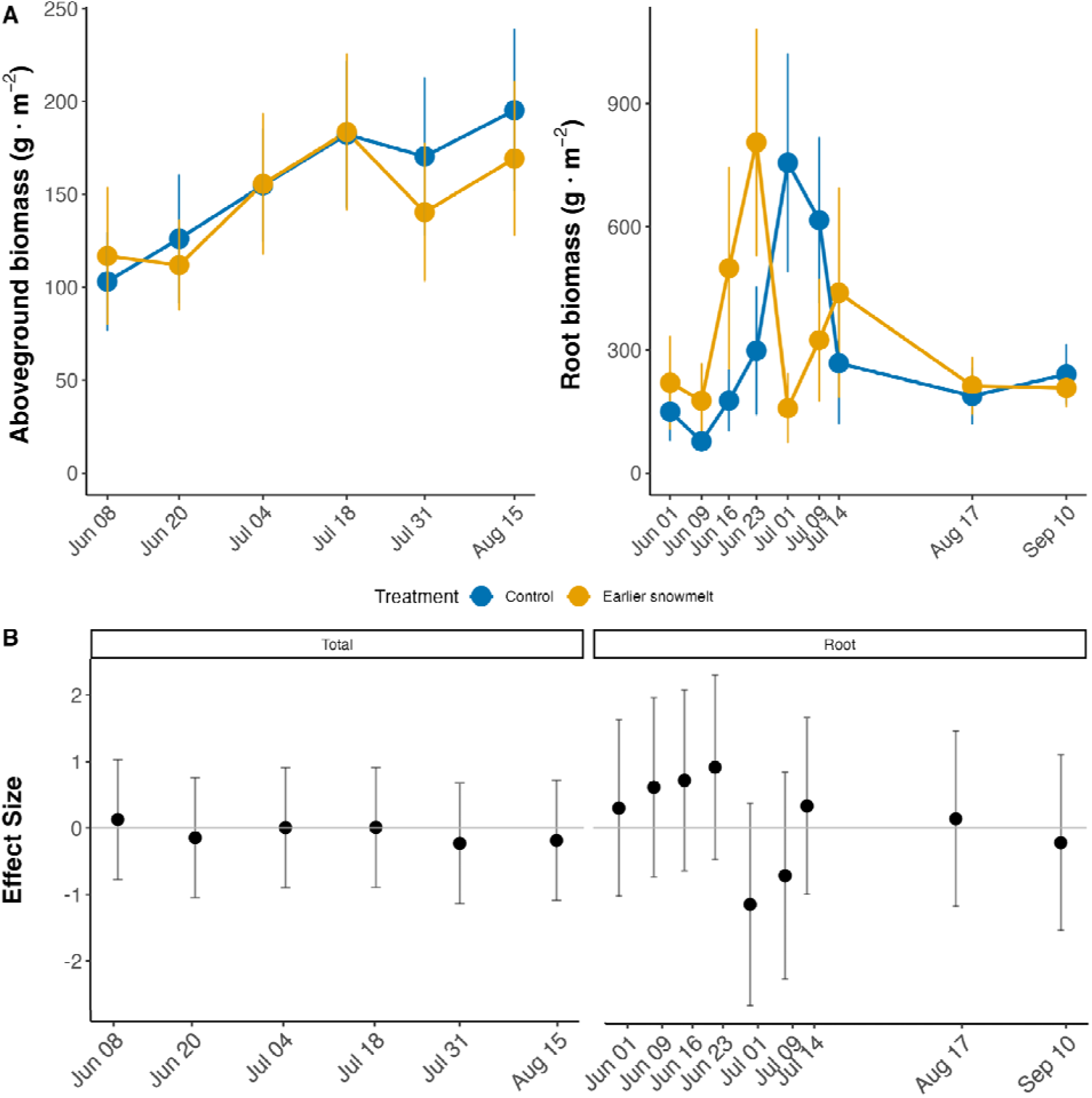
a) Aboveground biomass and root biomass over time in control and earlier snowmelt plots. The error bars represent the standard error. b) The effect sizes of the treatment on aboveground biomass and root biomass. The effect sizes are calculated using Hedge’s d, comparing the control versus the snowmelt plots. Above the zero line indicates that the biomass value in the snowmelt plots was greater than the control plots. The error bars represent the 95% confidence interval.

At the first sampling point in early June, graminoid biomass was 78% higher in the earlier snowmelt plots (Figure 3b), and had a significant effect size. On average, treatment plots had 47% more graminoid biomass throughout the growing season (Figure 3a). Graminoid biomass also changed significantly over the growing season (Table 1). In contrast to graminoids, shrub biomass was 22% lower in the earlier snowmelt plots over the growing season (Figure 3a), with the effect being more pronounced at the end of the season than at the beginning of the season (Figure 3b), though not statistically significant. Shrub biomass changed significantly over the growing season (Table 1), generally increasing throughout the season (Figure 3a). Forb biomass was not affected by earlier snowmelt but did show significant seasonal changes, peaking in mid-July and declining for the remainder of the growing season.

**Figure 3.**
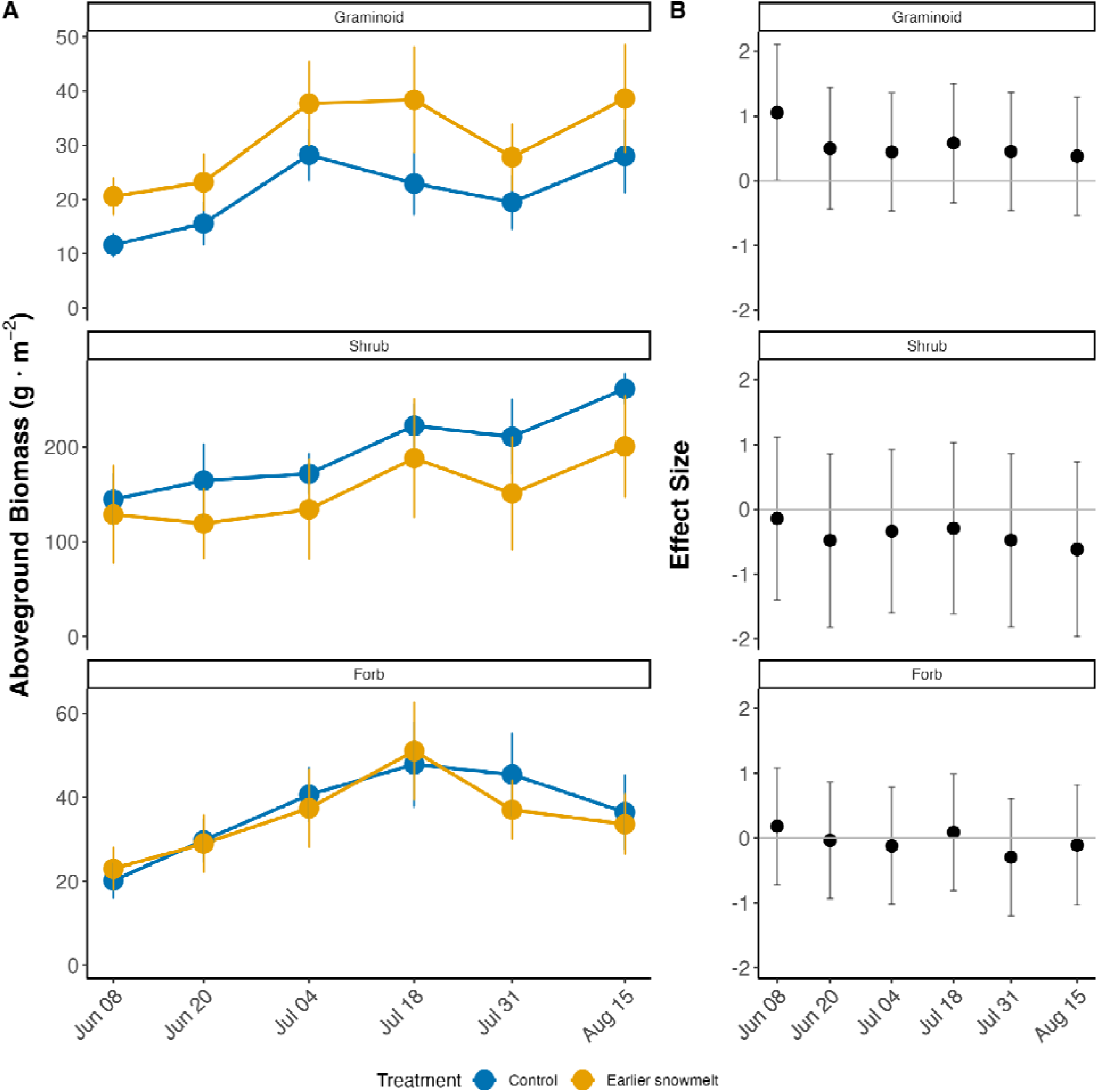
a) Graminoid, shrub, and forb biomass in control and earlier snowmelt plots over time. The error bars represent the standard error. b) The effect sizes of the treatment on aboveground biomass and root biomass. The effect sizes are calculated using Hedge’s d, comparing the control versus the earlier snowmelt plots. Above the zero line indicates that the biomass value in the earlier snowmelt plots was greater than the control plots. The error bars represent the 95% confidence interval.

NEE changed significantly over the season (Figure 4a, Table 1). Both the control and snowmelt plots were carbon sources in the spring but became carbon sinks by early July. At no point was there a significant difference in NEE between the earlier snowmelt and the control plots (Figure 4b). GPP also varied significantly over the growing season, starting low, peaking in early July, and then decreasing through the end of the season. There was a significant treatment × time interaction for GPP (Table 1). In early June, the advanced snowmelt plots showed 206% greater GPP than the control plots and had a significant effect size, but this effect diminished over the season, and by mid-July, GPP in the snowmelt plots was less than in the control plots (Figure 4b). ER did not change significantly over the season and was not significantly impacted by the treatment.

**Figure 4.**
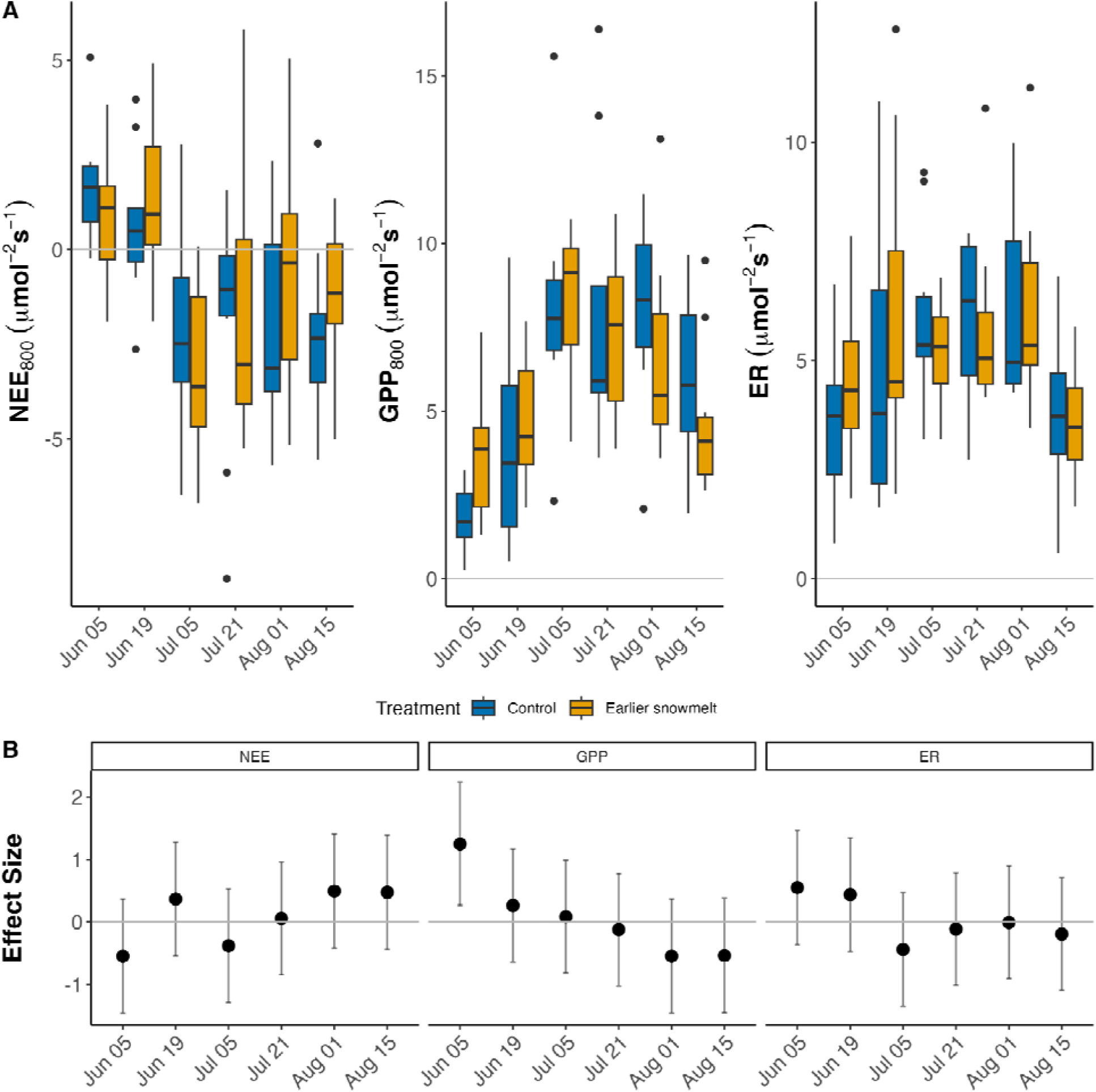
a) NEE, GPP, and ER across the growing season in earlier snowmelt and control plots. Carbon fluxes are on the y-axis, and the date is on the x-axis. For each box, the center line represents the median, and the box represents the interquartile range, with the bottom of the box the 25th percentile, and the top the 75th percentile. The whiskers extend to the largest and smallest data points within 1.5 times the interquartile range. b) The effect sizes of the treatment on NEE, GPP, and ER. The effect sizes are calculated using Hedge’s d, comparing the control versus the snowmelt plots. Values above the zero line indicate that the carbon value in the snowmelt plots was greater than the control plots. For NEE, any effect size values above the zero line indicate that earlier snowmelt plots are a greater carbon source. The error bars represent the 95% confidence interval.

The model fit for predicted NEE over the entire growing season (predicted vs. observed *r*^2^) was 0.68 (Figure S3). Light was a significant predictor of NEE in both the control and earlier snowmelt plots across the entire growing season, such that more light resulted in a larger carbon sink (Figure 5a). The strongest effect of light was during the mid-season, resulting in the strongest carbon sink for all plots. In the early season, the light parameter estimate for the earlier snowmelt plots was significantly different from the control plots, indicating a much stronger carbon sink in the earlier snowmelt plots. Neither air temperature nor soil moisture were a significant parameter at any time point (Figure 5a). However, increased soil moisture generally resulted in a stronger carbon sink, and increased air temperature resulted in a weaker carbon sink, with the exception being the control plots, which had an opposite relationship in the early season. NEE varied over the growing season and was a weak carbon sink in the early growing season, a strong carbon sink in the mid-season, and decreased again in the late season (Figure 5b). Over the entire growing season, cumulative NEE was 27.9 ± 17.7 mol CO_2_ m^-2^ in the control plots and 33.9 ± 18.1 mol CO_2_ m^-2^ in the earlier snowmelt plots, a 22% increase in carbon sink strength with earlier snowmelt (Figure 5c). Across the growing season, the earlier snowmelt plots were a 413% stronger carbon sink in the early season, 37% stronger carbon sink in the mid-season, and 33% weaker carbon sink in the late season.

**Figure 5.**
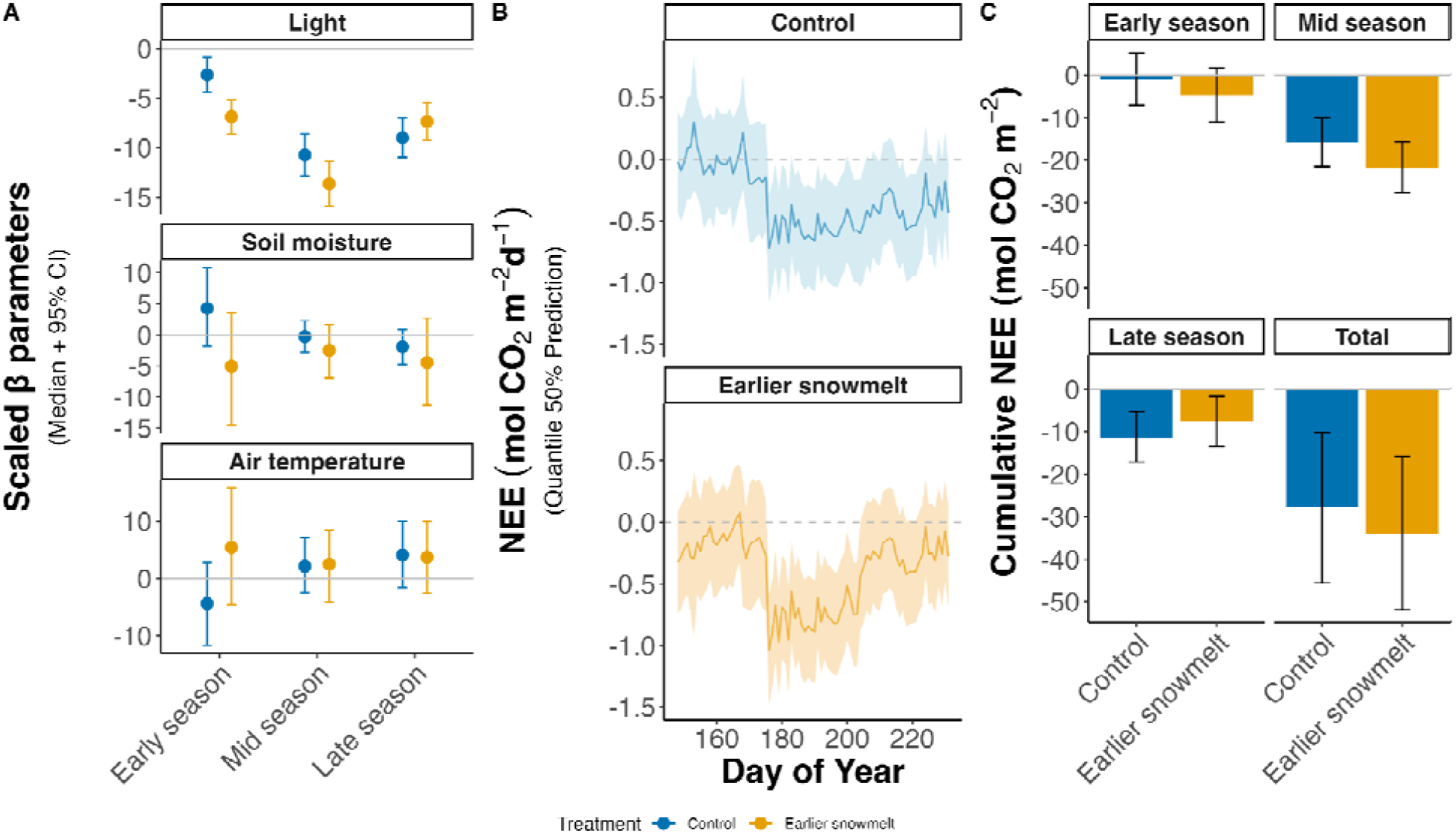
a) Scaled parameter values for the Bayesian model, median plus 95% confidence interval. Parameter values are scaled by the mean of the factor (i.e., the soil moisture parameter in the early season is multiplied by the mean soil moisture value in the early season). Values that do not cross zero are significant. Any number below zero indicates that the parameter causes a larger carbon sink, and any number above zero indicates that the parameter causes a greater carbon source. b) NEE from late May to late August in earlier snowmelt and control plots. Data were modeled using a Bayesian model, and NEE was predicted based on light, soil moisture, and air temperature values. The blue and orange lines are the 50% quantile measurements, and the light blue and light orange areas around are the 95% confidence interval. NEE values below the zero line indicate a carbon sink, while NEE values above the zero line indicate a carbon source. c) Cumulative NEE, summed from late May to late August in control and earlier snowmelt plots and for each season. Error bars represent standard deviation.

Forb biomass was a significant predictor of NEE, such that increased forb biomass resulted in lower NEE (*p* = 0.05, Table S3, Figure S4). Shrub biomass was also a significant predictor of NEE, with greater biomass resulting in lower NEE (*p* = 0.003, Table S3, Figure S4). Graminoid biomass interacted with time to have a marginally significant effect on NEE (*p* = 0.06, Table S3, Figure S4). Later in the season, graminoid biomass had a positive relationship with NEE, with more graminoid biomass resulting in larger NEE, while there was a slightly negative relationship in the early season (Figure S4). Graminoid biomass and time interacted to influence GPP, with the relationship becoming more positive later in the season (*p* = 0.02; Table S3, Figure S4).

Graminoid, shrub, and forb biomass were all negatively correlated with GPP, such that increases in biomass led to more negative GPP (*p* = 0.03, 0.04, and 0.001, respectively, Table S3, Figure S4). Only forb biomass had a significant relationship with ER, with greater forb biomass resulting in less ER (*p* = 0.005, Table S3, Figure S4).

## Discussion

Winters are changing, and earlier snowmelt is occurring more frequently with climate change (Gottlieb & Mankin, 2024). However, there is still uncertainty about how earlier snowmelt will impact the carbon balance of ecosystems. We advanced snowmelt by 12 days in a montane meadow and found that earlier snowmelt not only shifted the growing season earlier but also increased the strength of the carbon sink. Earlier snowmelt resulted in a stronger carbon sink at the start of the growing season and through peak season, and though it weakened the carbon sink at the end of the season, earlier snowmelt resulted in greater carbon accumulation (more negative NEE) across the entire season. The timing of aboveground green-up and root growth shifted earlier in the year due to earlier snowmelt. Shrubs and graminoids showed the strongest responses, with higher graminoid biomass in early snowmelt plots, especially in the early season, suggesting that graminoids’ plastic phenological response could increase their abundance in response to changing snowmelt date. Overall, our results show that relatively small shifts in snowmelt date could have large implications for carbon storage in montane meadows.

Total aboveground biomass did not change with an earlier snowmelt date, supporting our hypothesis and previous work suggesting that drought might have a larger effect on total aboveground biomass than snowmelt timing (Möhl et al., 2023). Contrary to our hypothesis, total root biomass remained unchanged with earlier snowmelt, despite contradictory findings in other studies. Some studies show that increased snow leads to more root biomass (Li et al., 2020), while others show the opposite (D’Imperio et al., 2018). Root biomass typically responds to soil moisture, increasing in drier ecosystems to optimize water acquisition (Liu et al., 2018).

However, roots can maintain similar biomass while changing branching patterns or their diameters in response to water stress (Yetgin, 2024). In our experiment, earlier snowmelt may not have significantly altered soil moisture enough to impact overall root biomass, especially given how similar soil moisture was between the treatments in the early season.

In contrast to total above or below-ground biomass, earlier snowmelt shifted root phenology and the timing of aboveground green-up. Peak root growth occurred earlier in the year, which contrasts with previous studies showing that root phenology is relatively unresponsive to snowmelt timing (Blume-Werry et al., 2017). A recent meta-analysis of 88 studies found that herbaceous root phenology was unaffected by warming, while aboveground herbaceous plants shifted their growing season earlier (Liu et al., 2022). However, warmer temperatures throughout the growing season and changes in the snowmelt timing could affect root phenology differently. Despite peak root biomass shifting earlier with advanced snowmelt, we did not observe a shift in the timing of aboveground peak biomass. However, earlier snowmelt did lead to earlier green-up (higher NDVI; Figure 1) and greater early productivity (higher GPP; Figure 4b). The disparity between the treatment effects on the timing of biomass production and green-up suggests that, though the biomass of perennial woody structures remained stable, green-up timing and productivity were more responsive to earlier snowmelt. Traditionally, phenology focuses on life history events such as bud burst and flowering (Gezon et al., 2016; Inouye, 2008; Sherwood et al., 2017; Slatyer et al., 2022). However, focusing on the timing of ecosystem functions, such as productivity (Chen et al., 2019; Galvagno et al., 2013), can clarify how changing snowmelt timing affects the carbon cycle on the ecosystem scale.

In plots with earlier snowmelt, graminoid biomass was higher, especially early in the season, and shrub biomass was lower throughout the season. Graminoids are particularly responsive to changes in snowmelt timing and winter snowpack, likely due to shifts in soil moisture (Ma et al., 2024). However, the response of graminoids to drought conditions can vary depending on the plant community context and the relative soil moisture of the ecosystem (Gherardi & Sala, 2015; P. Li et al., 2020; Liu et al., 2018; Rosbakh et al., 2017). Since grasses typically have relatively shallow fibrous roots, the early-season moisture pulse from snowmelt may have stimulated their growth, sustaining high biomass throughout the growing season. Previous work supports that early-season abiotic conditions have a large impact on total aboveground biomass throughout the growing season (Guo et al., 2017; Shaw et al., 2022). Additionally, snowmelt is often coupled with a pulse of nitrogen (Miller et al., 2009), and graminoids are more responsive to short-term nutrient changes than shrubs (Carbonell-Silletta et al., 2024). While shrubs tend to increase their biomass with warming temperatures (Amagai et al., 2018; Broadbent et al., 2022) and have strong phenological responses to changes in snowmelt timing (Wipf & Rixen, 2010), we found lower shrub biomass with earlier snowmelt. This could indicate that shrub biomass responses to changes in snowmelt date differ from warming effects, as shrubs tend to be slow-growing and could be relatively resilient to small changes in snowmelt date, and may respond more strongly to long-term warming. Changes in plant community composition can also affect the carbon balance, as graminoids, shrubs, and forbs process carbon differently across the growing season. Graminoids differed in their relationship between GPP and NEE at different points in the season, with graminoids tending to have more carbon uptake in the earlier season, and that trend reversed as the season progressed. Surprisingly, this trend was not detected statistically in forbs or shrubs. This suggests that graminoids could be more phenologically sensitive than other functional groups (Bektaş et al., 2021) and can advance their growing season, driving the difference in early-season GPP in the earlier snowmelt plots.

Our study found that early snowmelt had a greater effect on GPP than ER. This is similar to findings from 119 site-years of eddy flux tower data in the Arctic, where productivity was impacted by snowmelt timing, but there was little impact on ecosystem respiration (Zona et al., 2022). Similarly, a meta-analysis of 49 studies looking at the effect of snow depth on carbon dynamics found no substantial impact of snow depth on respiration (Li et al., 2016), consistent with our results. One study found that earlier snowmelt influences annual NEE by affecting respiration; however, this effect is attributed to reductions in winter respiration caused by earlier snowmelt rather than changes in growing season respiration (Galvagno et al., 2013). We observed a significant interaction between treatment and time for productivity but not for respiration. This finding is consistent with other studies showing that earlier snowmelt boosts early-season productivity (Chen et al., 2019; Sherwood et al., 2017; Vorkauf et al., 2021; Wipf & Rixen, 2010; Zona et al., 2022), although many studies focus more on phenology than on productivity (Wipf & Rixen, 2010). A review of snow manipulation experiments in Arctic and alpine ecosystems found that flower phenology was particularly responsive to the snowmelt timing and tracked earlier snowmelt (Wipf & Rixen, 2010), likely boosting early plant production. In our study, the increase in GPP with early snowmelt was only significant early in the season, and diminished as the season progressed.

We found that montane meadows with earlier snowmelt dates acted as stronger carbon sinks in the early and mid season. Though other work has also found that the growing season will be shifted forward, and not extended, with earlier snowmelt and climate change (Möhl et al., 2022; Zona et al., 2022), we also find an overall increase in the carbon sink due to increased uptake in both the early and mid season. Interestingly, we found that light was the strongest predictor of NEE, more than air temperature or soil moisture, which could support the idea of periodicity, or that plants will be cued by light with a changing snowmelt date but maintain a fixed growing period, leading to earlier senescence (Semenchuk et al., 2016). Most studies examining carbon fluxes in relation to snow changes use natural variation in snowmelt days across years (Chen et al., 2019; Galvagno et al., 2013; López-Blanco et al., 2017; Zona et al., 2022), with varying results. Some studies report that earlier snowmelt shifts the growing season forward, increasing productivity in the spring (Chen et al., 2019; Zona et al., 2022), while others find no effect on NEE because changes in GPP and ER cancel each other out (López-Blanco et al., 2017). In some cases, earlier snowmelt decreased NEE due to lower respiration rates (Galvagno et al., 2013).

Our study is one of the first to measure NEE in response to experimentally manipulated snowmelt timing. Our results indicate that earlier snowmelt in montane meadows will cause productivity to not only shift earlier but also to amplify.

Environmental gradients alone can not fully capture climate change effects as they do not have any controls, making experiments valuable for new insights as they can pinpoint mechanisms while accounting for confounding factors (Metz & Tielbörger, 2016; Rysavy et al., 2014).

Integrating natural observations and experiments is key to understanding climate change impacts (Prager et al., 2022). Approximately 55% of snow studies rely on observations and 45% on experiments, but most focus on plant phenology or community diversity responses (Slatyer et al., 2022). Natural gradients usually exhibit larger variations in snowmelt timing than experiments, potentially leading to greater ecosystem responses (Rixen et al., 2022). However, natural gradients tend to incorporate areas with frequent early snowmelt due to their slope or aspect, where vegetation is likely adapted to variable conditions (Rixen et al., 2022). Ecosystem manipulation experiments offer a way to assess the impacts of earlier snowmelt on ecosystems where snowmelt variation is historically minimal, which is important as climate change alters ecosystems with historically less variation. However, experiments also have limitations, as they often cover less spatial and temporal variability. For example, 2023 had a large snowpack and a relatively dry growing season (Figure S2b), potentially amplifying the effects of an earlier snowmelt. In a year with a smaller snowpack and greater summer precipitation, the results could be less pronounced.

In conclusion, earlier snowmelt strengthens the carbon sink strength in montane meadows by advancing the growing season and increasing carbon uptake in the midseason. Graminoid biomass was higher in response to earlier snowmelt, driving the early-season productivity increases. Phenological shifts in plants both above-and belowground will occur with earlier snowmelt. In our experimental meadow, snowmelt has advanced by 3.5 days per decade (Iler et al., 2013), and we advanced snowmelt by about 12 days. Our results, therefore, estimate the carbon balance approximately three decades into the future, around 2055. Our study underscores how even small environmental changes, such as a less-than-two-week advance in snowmelt, can significantly affect ecosystems, potentially increasing carbon storage in montane meadows.

Understanding how ecosystems respond to changes in snowmelt timing is crucial for predicting the impacts of climate variability.

## Notes

**Acknowledgments** This study was supported by the U.S. Department of Energy, Office of Science, Office of Biological and Environmental Research, Terrestrial Ecosystem Sciences program under award number DE-FOA-0002392. PS was partially supported by the Watershed Function Scientific Focus Area at Lawrence Berkeley National Laboratory, funded by the U.S. Department of Energy, Office of Science, Office of Biological and Environmental Research under Award Number DE-AC02-05CH11231. NDVI data were provided by Ian Breckheimer and Amanda Henderson at Rocky Mountain Biological Laboratory. This project was funded by a Rocky Mountain Biological Laboratory Graduate Student Fellowship, an Institute of Global Change Biology Fellowship, and the Dr. Nancy Williams Walls Award for Field Research to OKV.

### Competing Interest Statement

The authors have declared no competing interest.

### Summary of Updates

The cumulative NEE model has been revised, adding light as a parameter and estimating the parameters three times across the growing season. We now see that earlier snowmelt increased the carbon sink strength by 22%, with the strongest effect in the early spring. The effect diminished as the growing season progressed, and the control plots were a greater carbon sink in the later season.

## References

Amagai, Y., Kudo, G., & Sato, K. (2018). Changes in alpine plant communities under climate change: Dynamics of snow-meadow vegetation in northern Japan over the last 40 years. Applied Vegetation Science, 21(4), 561–571. 10.1111/AVSC.12387

Bektaş, B., Thuiller, W., Saillard, A., Choler, P., Renaud, J., Colace, M.-P., Della Vedova, R., & Münkemüller, T. (2021). Lags in phenological acclimation of mountain grasslands after recent warming. Journal of Ecology, 109(9), 3396–3410. 10.1111/1365-2745.13727

Blume-Werry, G., Jansson, R., & Milbau, A. (2017). Root phenology unresponsive to earlier snowmelt despite advanced above-ground phenology in two subarctic plant communities. Functional Ecology, 31(7), 1493–1502. 10.1111/1365-2435.12853

Broadbent, A. A. D., Bahn, M., Pritchard, W. J., Newbold, L. K., Goodall, T., Guinta, A., Snell, H. S. K., Cordero, I., Michas, A., Grant, H. K., Soto, D. X., Kaufmann, R., Schloter, M., Griffiths, R. I., & Bardgett, R. D. (2022). Shrub expansion modulates belowground impacts of changing snow conditions in alpine grasslands. Ecology Letters, 25(1), 52–64. 10.1111/ELE.13903

Carbonell-Silletta, L., Scholz, F. G., Burek, A., Villa, V. D., Cavallaro, A., Askenazi, J. O., Arias, N. S., Hao, G.-Y., Goldstein, G., & Bucci, S. J. (2024). Nitrogen rather than water availability limits aboveground primary productivity in an arid ecosystem: Substantial differences between grasses and shrubs. Ecohydrology, 17(3), e2636. 10.1002/eco.2636

Chen, S., Huang, Y., & Wang, G. (2019). Response of vegetation carbon uptake to snow-induced phenological and physiological changes across temperate China. Science of The Total Environment, 692, 188–200. 10.1016/j.scitotenv.2019.07.222

Darrouzet-Nardi, A., Steltzer, H., Sullivan, P. F., Segal, A., Koltz, A. M., Livensperger, C., Schimel, J. P., & Weintraub, M. N. (2019). Limited effects of early snowmelt on plants, decomposers, and soil nutrients in Arctic tundra soils. Ecology and Evolution, 9(4), 1820–1844. 10.1002/ECE3.4870

D’Imperio, L., Arndal, M. F., Nielsen, C. S., Elberling, B., & Schmidt, I. K. (2018). Fast responses of root dynamics to increased snow deposition and summer air temperature in an arctic wetland. Frontiers in Plant Science, 9, 1258. 10.3389/FPLS.2018.01258/BIBTEX

Galvagno, M., Wohlfahrt, G., Cremonese, E., Rossini, M., Colombo, R., Filippa, G., Julitta, T., Manca, G., Siniscalco, C., Morra Di Cella, U., & Migliavacca, M. (2013). Phenology and carbon dioxide source/sink strength of a subalpine grassland in response to an exceptionally short snow season. Environmental Research Letters, 8(2), 025008. 10.1088/1748-9326/8/2/025008

Gamon, J. A., Huemmrich, K. F., Stone, R. S., & Tweedie, C. E. (2013). Spatial and temporal variation in primary productivity (NDVI) of coastal Alaskan tundra: Decreased vegetation growth following earlier snowmelt. Remote Sensing of Environment, 129, 144–153. 10.1016/j.rse.2012.10.030

Ganjurjav, H., Gao, Q., Schwartz, M. W., Zhu, W., Liang, Y., Li, Y., Wan, Y., Cao, X., Williamson, M. A., Jiangcun, W., Guo, H., & Lin, E. (2016). Complex responses of spring vegetation growth to climate in a moisture-limited alpine meadow. Scientific Reports 2016 6:1, 6(1), 1–10. 10.1038/srep23356

Gezon, Z. J., Inouye, D. W., & Irwin, R. E. (2016). Phenological change in a spring ephemeral: Implications for pollination and plant reproduction. Global Change Biology, 22(5), 1779–1793. 10.1111/GCB.13209

Gherardi, L. A., & Sala, O. E. (2015). Enhanced precipitation variability decreases grass-and increases shrub-productivity. Proceedings of the National Academy of Sciences, 112(41), 12735–12740. 10.1073/pnas.1506433112

Gottlieb, A. R., & Mankin, J. S. (2024). Evidence of human influence on Northern Hemisphere snow loss. Nature, 625(7994), 293–300. 10.1038/s41586-023-06794-y

Guo, L., Cheng, J., Luedeling, E., Koerner, S. E., He, J. S., Xu, J., Gang, C., Li, W., Luo, R., & Peng, C. (2017). Critical climate periods for grassland productivity on China’s Loess Plateau. Agricultural and Forest Meteorology, 233, 101–109. 10.1016/J.AGRFORMET.2016.11.006

Hu, J., Hopping, K. A., Bump, J. K., Kang, S., & Klein, J. A. (2013). Climate Change and Water Use Partitioning by Different Plant Functional Groups in a Grassland on the Tibetan Plateau. PLOS ONE, 8(9), e75503. 10.1371/JOURNAL.PONE.0075503

Hubbard, S. S., Williams, K. H., Agarwal, D., Banfield, J., Beller, H., Bouskill, N., Brodie, E., Carroll, R., Dafflon, B., Dwivedi, D., Falco, N., Faybishenko, B., Maxwell, R., Nico, P., Steefel, C., Steltzer, H., Tokunaga, T., Tran, P. A., Wainwright, H., & Varadharajan, C. (2018). The East River, Colorado, Watershed: A Mountainous Community Testbed for Improving Predictive Understanding of Multiscale Hydrological–Biogeochemical Dynamics. Vadose Zone Journal, 17(1), 180061. 10.2136/vzj2018.03.0061

Ibanez, M., Altimir, N., Ribas, A., Eugster, W., & Sebastia, M. T. (2020). Phenology and plant functional type dominance drive CO2 exchange in seminatural grasslands in the Pyrenees. The Journal of Agricultural Science, 158(1–2), 3–14. 10.1017/S0021859620000179

Iler, A. M., Høye, T. T., Inouye, D. W., & Schmidt, N. M. (2013). Nonlinear flowering responses to climate: Are species approaching their limits of phenological change? Philosophical Transactions of the Royal Society B: Biological Sciences, 368(1624), 20120489. 10.1098/rstb.2012.0489

Inouye, D. W. (2008). Effects of climate change on phenology, frost damage, and floral abundance of montane wildflowers. Ecology, 89(2), 353–362.

Kawai, Y., & Kudo, G. (2021). Climate change shifts population structure and demographics of an alpine herb, Anemone narcissiflora ssp. Sachalinensis (Ranunculaceae), along a snowmelt gradient. Population Ecology, 63(3), 260–271. 10.1002/1438-390X.12089

Kelsey, K. C., Pedersen, S. H., Leffler, A. J., Sexton, J. O., Feng, M., & Welker, J. M. (2021). Winter snow and spring temperature have differential effects on vegetation phenology and productivity across Arctic plant communities. Global Change Biology, 27(8), 1572– 1586. 10.1111/GCB.15505

Kudo, G. (2021). Habitat-specific effects of flowering advance on fruit-set success of alpine plants: A long-term record of flowering phenology and fruit-set success of Rhododendron aureum. Alpine Botany, 131(1), 53–62. 10.1007/S00035-021-00248-9/FIGURES/6

Lambert, A. M., Miller-Rushing, A. J., & Inouye, D. W. (2010). Changes in snowmelt date and summer precipitation affect the flowering phenology of Erythronium grandiflorum (glacier lily; Liliaceae). American Journal of Botany, 97(9), 1431–1437. 10.3732/AJB.1000095

Li, P., Sayer, E. J., Jia, Z., Liu, W., Wu, Y., Yang, S., Wang, C., Yang, L., Chen, D., Bai, Y., & Liu, L. (2020). Deepened winter snow cover enhances net ecosystem exchange and stabilizes plant community composition and productivity in a temperate grassland. Global Change Biology, 26(5), 3015–3027. 10.1111/GCB.15051

Li, W., Wu, J., Bai, E., Jin, C., Wang, A., Yuan, F., & Guan, D. (2016). Response of terrestrial carbon dynamics to snow cover change: A meta-analysis of experimental manipulation (II). Soil Biology and Biochemistry, 103, 388–393. 10.1016/j.soilbio.2016.09.017

Liu, H., Mi, Z., Lin, L., Wang, Y., Zhang, Z., Zhang, F., Wang, H., Liu, L., Zhu, B., Cao, G., Zhao, X., Sanders, N. J., Classen, A. T., Reich, P. B., & He, J.-S. (2018). Shifting plant species composition in response to climate change stabilizes grassland primary production. Proceedings of the National Academy of Sciences, 115(16), 4051–4056. 10.1073/pnas.1700299114

Liu, H., Wang, H., Li, N., Shao, J., Zhou, X., van Groenigen, K. J., & Thakur, M. P. (2022). Phenological mismatches between above-and belowground plant responses to climate warming. Nature Climate Change, 12(1), 97–102. 10.1038/s41558-021-01244-x

Liu, H., Xiao, P., Zhang, X., Chen, S., Wang, Y., & Wang, W. (2023). Winter snow cover influences growing-season vegetation productivity non-uniformly in the Northern Hemisphere. Communications Earth & Environment, 4(1), 1–10. 10.1038/s43247-023-01167-9

López-Blanco, E., Lund, M., Williams, M., Tamstorf, M. P., Westergaard-Nielsen, A., Exbrayat, J. F., Hansen, B. U., & Christensen, T. R. (2017). Exchange of CO2 in Arctic tundra: Impacts of meteorological variations and biological disturbance. Biogeosciences, 14(19), 4467–4483. 10.5194/bg-14-4467-2017

Ma, W., Hu, J., Zhang, B., Guo, J., Zhang, X., & Wang, Z. (2024). Later-melting rather than thickening of snowpack enhance the productivity and alter the community composition of temperate grassland. Science of The Total Environment, 923, 171440. 10.1016/j.scitotenv.2024.171440

McLaren, J. R., & Turkington, R. (2011). Biomass compensation and plant responses to 7 years of plant functional group removals. Journal of Vegetation Science, 22(3), 503–515. 10.1111/j.1654-1103.2011.01263.x

Metz, J., & Tielbörger, K. (2016). Spatial and temporal aridity gradients provide poor proxies for plant–plant interactions under climate change: A large-scale experiment. Functional Ecology, 30(1), 20–29. 10.1111/1365-2435.12599

Miller, A. E., Schimel, J. P., Sickman, J. O., Skeen, K., Meixner, T., & Melack, J. M. (2009). Seasonal variation in nitrogen uptake and turnover in two high-elevation soils: Mineralization responses are site-dependent. Biogeochemistry, 93(3), 253–270. 10.1007/S10533-009-9301-4/TABLES/4

Möhl, P., von Büren, R. S., & Hiltbrunner, E. (2022). Growth of alpine grassland will start and stop earlier under climate warming. Nature Communications, 13(1), 7398. 10.1038/s41467-022-35194-5

Möhl, P., Vorkauf, M., Kahmen, A., & Hiltbrunner, E. (2023). Recurrent summer drought affects biomass production and community composition independently of snowmelt manipulation in alpine grassland. Journal of Ecology, 111(11), 2357–2375. 10.1111/1365-2745.14180

Pepin, N., Bradley, R. S., Diaz, H. F., Baraer, M., Caceres, E. B., Forsythe, N., Fowler, H., Greenwood, G., Hashmi, M. Z., Liu, X. D., Miller, J. R., Ning, L., Ohmura, A., Palazzi, E., Rangwala, I., Schöner, W., Severskiy, I., Shahgedanova, M., Wang, M. B.,… Yang, D. Q. (2015). Elevation-dependent warming in mountain regions of the world. Nature Climate Change 2015 5:5, 5(5), 424–430. 10.1038/NCLIMATE2563

Pinheiro, J. C., & Bates, D. M. (2000). Mixed-Effects Models in S and S-PLUS. Springer. 10.1007/b98882

Pinheiro, J. C., Bates, D., & R Core Team. (2023). nlme: Linear and Nonlinear Mixed Effects Models (R package version 3.1-163). https://CRAN.R-project.org/package=nlme

Plummer, M. (2003). JAGS: A Program for Analysis of Bayesian Graphical Models Using Gibbs Sampling. Proceedings of the 3rd International Workshop on Distributed Statistical Computing (DSC 2003), 1–10.

Plummer, M. (2023). rjags: Bayesian Graphical Models using MCMC (R package version 4-15). https://CRAN.R-project.org/package=rjags

Prager, C. M., Classen, A. T., Sundqvist, M. K., Barrios-Garcia, M. N., Cameron, E. K., Chen, L., Chisholm, C., Crowther, T. W., Deslippe, J. R., Grigulis, K., He, J.-S., Henning, J. A., Hovenden, M., Høye, T. T. T., Jing, X., Lavorel, S., McLaren, J. R., Metcalfe, D. B., Newman, G. S.,… Sanders, N. J. (2022). Integrating natural gradients, experiments, and statistical modeling in a distributed network experiment: An example from the WaRM Network. Ecology and Evolution, 12(10), e9396. 10.1002/ECE3.9396

Prager, C. M., Jing, X., Henning, J. A., Read, Q. D., Meidl, P., Lavorel, S., Sanders, N. J., Sundqvist, M., Wardle, D. A., Classen, A. T., Prager, C.:, Jing, X., Henning, J. A., Read, Q. D., Meidl, P., Lavorel, S., Sanders, N. J., Sundqvist, M., Wardle, D. A., & Classen, A. T. (2021). Climate and multiple dimensions of plant diversity regulate ecosystem carbon exchange along an elevational gradient. Ecosphere, 12(4). 10.1002/ecs2.3472

R Core Team. (2023). R: A Language and Environment for Statistical Computing. R Foundation for Statistical Computing. https://www.R-project.org/

Rixen, C., Thomas Høye, T., Macek, P., Aerts, R., Alatalo, J. M., Anderson, J. T., Arnold, P. A., Barrio, I. C., Bjerke, J. W., Björkman, M. P., Blok, D., Blume-Werry, G., Boike, J., Bokhorst, S., Carbognani, M., Christiansen, C. T., Convey, P., Cooper, E. J., Hans Cornelissen, J. C.,… Blok, D. (2022). Winters are changing: Snow effects on Arctic and alpine tundra ecosystems. 10.1139/as-2020-0058, 1–37. https://doi.org/10.1139/AS-2020-0058

Rosbakh, S., Leingärtner, A., Hoiss, B., Krauss, J., Steffan-Dewenter, I., & Poschlod, P. (2017). Contrasting Effects of Extreme Drought and Snowmelt Patterns on Mountain Plants along an Elevation Gradient. Frontiers in Plant Science, 8. 10.3389/fpls.2017.01478

Rysavy, A., Seifan, M., Sternberg, M., & Tielbörger, K. (2014). Shrub seedling survival under climate change – Comparing natural and experimental rainfall gradients. Journal of Arid Environments, 111, 14–21. 10.1016/j.jaridenv.2014.07.004

Sager, J., & McFarlane, J. (1997). Radiation. In Growth Chamber Handbook. https://www.controlledenvironments.org/wp-content/uploads/sites/6/2017/06/Ch01.pdf

Semenchuk, P. R., Gillespie, M. A. K., Rumpf, S. B., Baggesen, N., Elberling, B., & Cooper, E. J. (2016). High Arctic plant phenology is determined by snowmelt patterns but duration of phenological periods is fixed: An example of periodicity. Environmental Research Letters, 11(12), 125006. 10.1088/1748-9326/11/12/125006

Sengupta, M., Xie, Y., Lopez, A., Habte, A., Maclaurin, G., & Shelby, J. (2018). The National Solar Radiation Data Base (NSRDB). Renewable and Sustainable Energy Reviews, 89, 51–60. 10.1016/j.rser.2018.03.003

Shaw, E. A., White, C. T., Silver, W. L., Suding, K. N., & Hallett, L. M. (2022). Intra-annual precipitation effects on annual grassland productivity and phenology are moderated by community responses. Journal of Ecology, 110(1), 162–172. 10.1111/1365-2745.13792

Sherwood, J. A., Debinski, D. M., Caragea, P. C., & Germino, M. J. (2017). Effects of experimentally reduced snowpack and passive warming on montane meadow plant phenology and floral resources. Ecosphere, 8(3), e01745. 10.1002/ECS2.1745

Slatyer, R. A., Umbers, K. D. L., & Arnold, P. A. (2022). Ecological responses to variation in seasonal snow cover. Conservation Biology, 36(1), e13727. 10.1111/cobi.13727

Sundqvist, M. K., Sanders, N. J., Dorrepaal, E., Lindén, E., Metcalfe, D. B., Newman, G. S., Olofsson, J., Wardle, D. A., & Classen, A. T. (2020). Responses of tundra plant community carbon flux to experimental warming, dominant species removal and elevation. Functional Ecology, 34(7), 1497–1506. 10.1111/1365-2435.13567

Suzuki, R. O. (2014). Combined effects of warming, snowmelt timing, and soil disturbance on vegetative development in a grassland community. Plant Ecology, 215(12), 1399–1408. 10.1007/s11258-014-0396-x

Torchiano, M. (2020). effsize: Efficient Effect Size Computation (R package version 0.8.1). https://CRAN.R-project.org/package=effsize

Vorkauf, M., Kahmen, A., Körner, C., & Hiltbrunner, E. (2021). Flowering phenology in alpine grassland strongly responds to shifts in snowmelt but weakly to summer drought. Alpine Botany 2021 131:1, 131(1), 73–88. 10.1007/S00035-021-00252-Z

Wild, J., Kopecký, M., Macek, M., Šanda, M., Jankovec, J., & Haase, T. (2019). Climate at ecologically relevant scales: A new temperature and soil moisture logger for long-term microclimate measurement. Agricultural and Forest Meteorology, 268, 40–47. 10.1016/J.AGRFORMET.2018.12.018

Wipf, S., & Rixen, C. (2010). A review of snow manipulation experiments in Arctic and alpine tundra ecosystems. Polar Research, 29(1), Article 1. 10.3402/polar.v29i1.6054

Yan, S., Yin, L., Dijkstra, F. A., Wang, P., & Cheng, W. (2023). Priming effect on soil carbon decomposition by root exudate surrogates: A meta-analysis. Soil Biology and Biochemistry, 178, 108955. 10.1016/j.soilbio.2023.108955

Yetgin, A. (2024). Exploring the dynamic nature of root plasticity and morphology in the face of changing environments. Ecological Frontiers, 44(1), 112–119. 10.1016/j.chnaes.2023.07.008

Zhang, Q., Zhang, X., Lara, M. J., Li, Z., Xiao, J., Zhao, K., & Hu, T. (2023). Impacts of abiotic and biotic factors on tundra productivity near Utqiaġvik, Alaska. Environmental Research Letters, 18(9), 094070. 10.1088/1748-9326/acf7d6

Zona, D., Lafleur, P. M., Hufkens, K., Bailey, B., Gioli, B., Burba, G., Goodrich, J. P., Liljedahl, A. K., Euskirchen, E. S., Watts, J. D., Farina, M., Kimball, J. S., Heimann, M., Göckede, M., Pallandt, M., Christensen, T. R., Mastepanov, M., López-Blanco, E., Jackowicz-Korczynski, M.,… Oechel, W. C. (2022). Earlier snowmelt may lead to late season declines in plant productivity and carbon sequestration in Arctic tundra ecosystems. Scientific Reports 2022 12:1, 12(1), 1–10. 10.1038/s41598-022-07561-1

