## Supplemental Material for "Earlier snowmelt increases the strength of the carbon sink in montane meadows unequally across the growing season"

**Figures**

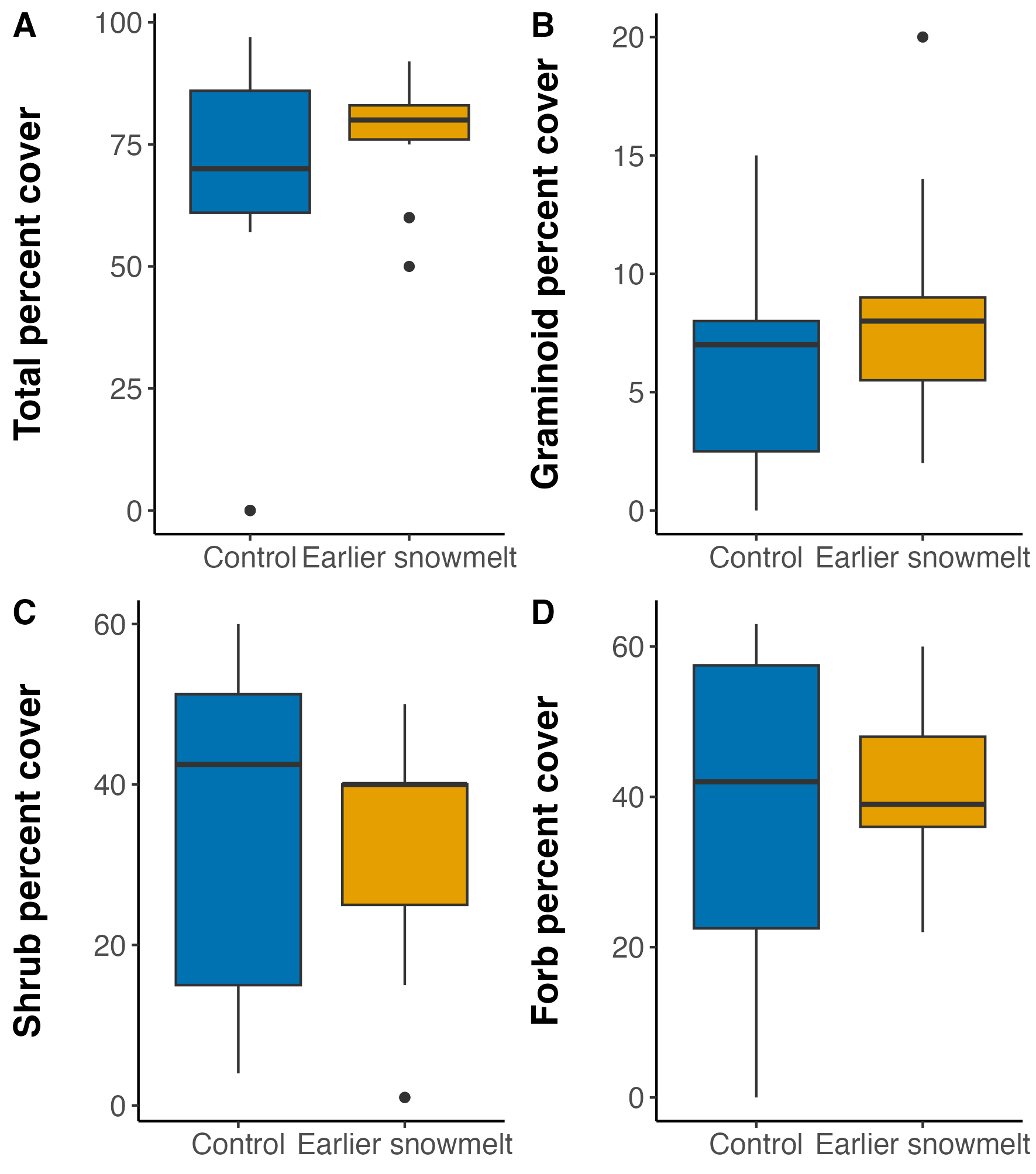

**Figure S1.** Pre-treatment (summer 2022) plant percent cover data from the plots. Notably, there is no significant difference between the plots pre-treatment.

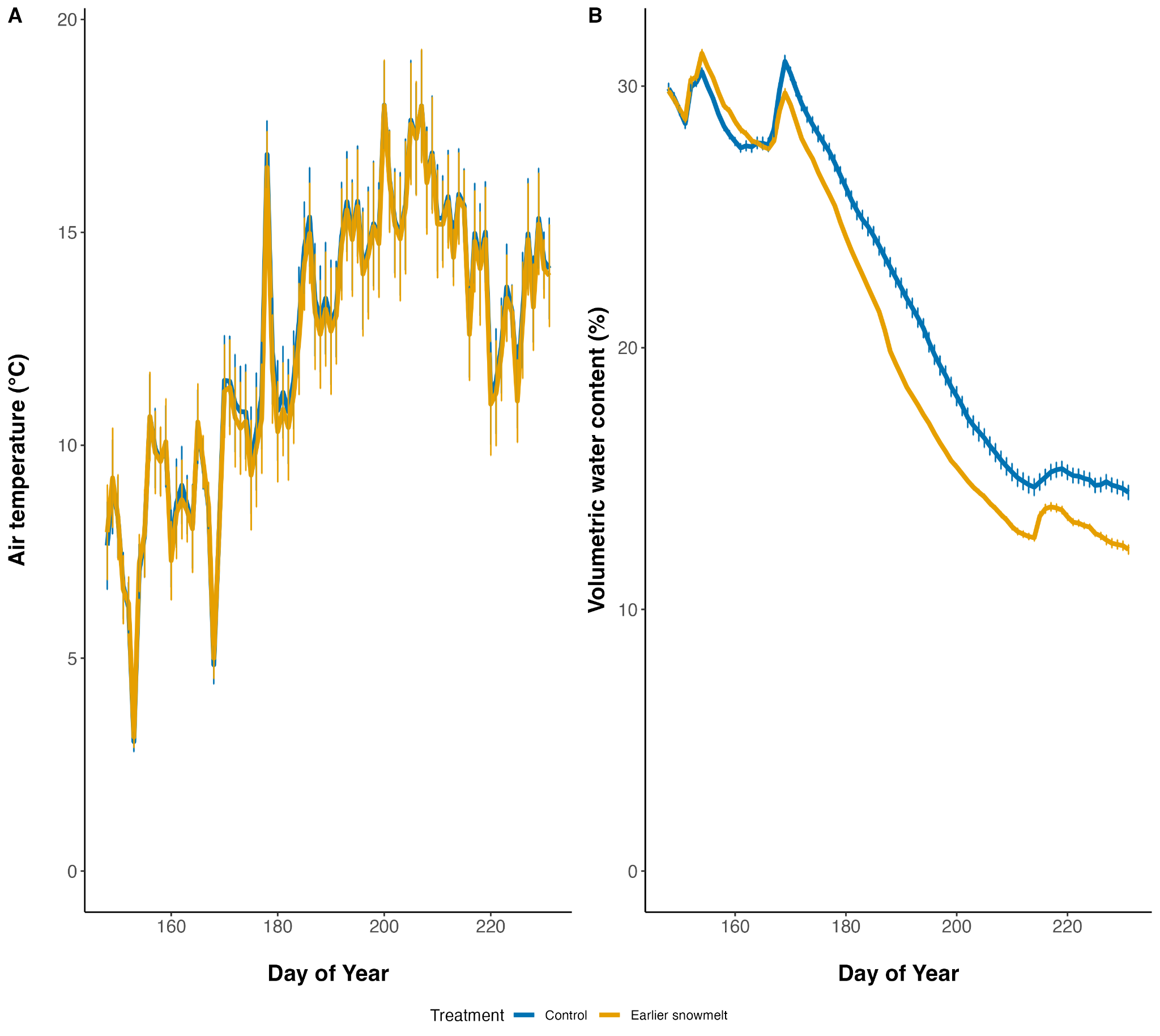

**Figure S2**. a) Average daily air temperature (°C) from late May to mid-August. b) Average daily volumetric water content (%) from late May to mid-August. Snowmelt occurred on day 136 of the year (May 16th) in the advanced snowmelt plots and day 148 of the year (May 28th) in the control plots.

Observed

| Predicted | NEE_even_season |
| --- | --- |
| 1.23139 | 0.624561 |
| 2.51267 | 1.63127 |
| 2.8457 | 1.88513 |
| 3.14261 | 3.28667 |
| -0.991578 | -1.53224 |
| 0.743312 | 0.985118 |
| 2.4318 | 1.38674 |
| 3.8664 | 4.68676 |
| 0.21154 | 1.91472 |
| 1.68023 | 1.98054 |
| 2.63168 | 1.42506 |
| 3.418 | 2.82455 |
| -4.926 | 0.183462 |
| 0.188711 | 0.248554 |
| 3.06089 | 0.72452 |
| 4.7344 | 4.41573 |
| 0.580892 | 0.210725 |
| 1.48773 | 1.27472 |
| 2.05249 | 0.938179 |
| 2.55933 | 1.99107 |
| -2.16691 | -4.74639 |
| 2.58724 | -0.727908 |
| 3.4411 | 2.10532 |
| 4.75272 | 4.85439 |
| -0.86729 | -1.55133 |
| 0.169051 | 0.200108 |
| 0.923152 | 0.791288 |
| 1.72744 | 0.170483 |
| -1.26256 | -1.04681 |
| 0.618185 | 3.13117 |
| 2.75785 | 3.10332 |
| 4.37692 | 6.26742 |
| -0.149178 | 1.11477 |
| 2.27872 | 0.373749 |
| 2.76945 | 5.38738 |
| -0.288333 | -1.11121 |
| 2.35988 | 2.99731 |
| 4.41402 | 2.03366 |
| 5.31998 | 5.24754 |
| 0.00215298 | -2.46373 |
| 1.35431 | 0.754529 |
| 1.81459 | 1.62222 |
| 2.37107 | 6.47262 |
| -2.27137 | -5.67583 |
| 1.23257 | -2.29981 |
| 2.77812 | 0.767339 |
| 4.52381 | 5.57569 |
| 1.18399 | -2.11555 |
| 1.90993 | 0.329749 |
| 2.56518 | 1.43554 |
| 3.09742 | 2.32805 |
| -3.99033 | -3.71519 |
| 0.0750163 | 0.440112 |
| 1.18449 | 1.18945 |
| 4.09173 | 4.35418 |
| -0.985909 | -0.503903 |
| 0.472361 | -0.157392 |
| 1.26709 | 0.33215 |
| 2.41929 | 2.85432 |
| -3.7171 | -6.99274 |
| -0.788706 | -1.71189 |
| 2.51142 | 1.28445 |
| 4.56859 | 8.94837 |
| -0.649459 | 0.525751 |
| 0.488887 | 0.9489 |
| 1.79076 | 1.21753 |
| 2.5266 | 1.67803 |
| -3.84193 | -1.59199 |
| 0.03253 | 1.05844 |
| 3.48006 | 2.7466 |
| 4.43983 | 4.70156 |
| -0.122745 | 1.24097 |
| 1.03962 | 2.15182 |
| 1.72936 | 4.89555 |
| 2.76887 | 8.07354 |
| -2.66704 | -0.617402 |
| 0.940227 | 2.27121 |
| 2.84992 | 4.45197 |
| 4.94505 | 8.32585 |
| -3.51331 | -5.03308 |
| 0.433317 | -1.13203 |
| 2.07984 | 0.927102 |
| 4.18642 | 6.50862 |
| -7.6943 | -9.61724 |
| -0.89402 | -1.95709 |
| 1.52591 | 1.47782 |
| 4.36796 | 7.65994 |
| -6.83084 | -10.6821 |
| -2.04562 | -2.96987 |
| 1.24205 | 0.428689 |
| 3.96376 | 6.96295 |
| -5.00384 | -4.62398 |
| -1.49782 | -0.00276291 |
| 1.75558 | 1.09973 |
| 4.17558 | 6.64213 |
| -9.22464 | -2.54216 |
| -4.25935 | -0.110329 |
| 2.08309 | 0.343604 |
| 3.94991 | 4.85088 |
| -4.5465 | -9.08745 |
| 0.730527 | -3.44399 |
| 2.16206 | -0.635659 |
| 4.22091 | 4.56174 |
| -3.94047 | -5.94873 |
| 0.534864 | -0.294838 |
| 2.75568 | 1.9314 |
| 3.98677 | 4.71772 |
| -5.63826 | -8.00814 |
| -0.477304 | -0.328792 |
| 1.92596 | 1.40468 |
| 3.65438 | 6.45444 |
| -2.79913 | -6.51757 |
| 0.725488 | -0.71123 |
| 2.33129 | 1.21264 |
| 3.97435 | 6.21706 |
| -8.28954 | -5.96302 |
| -2.04301 | -1.21501 |
| 0.593248 | 0.754719 |
| 4.44702 | 5.67873 |
| -3.31929 | -3.54167 |
| 0.831568 | -1.51842 |
| 2.58323 | 1.14884 |
| 4.59429 | 6.14895 |
| -6.9058 | -1.22627 |
| -0.091446 | 0.594805 |
| 2.58117 | 1.77266 |
| 5.26706 | 9.07387 |
| -5.27383 | -1.62676 |
| -2.18716 | -1.92971 |
| 2.01453 | -1.39728 |
| 4.30468 | 6.32396 |
| -4.93739 | -9.37221 |
| -0.265604 | -3.30513 |
| 3.14837 | 0.650737 |
| 5.29629 | 5.17264 |
| -4.39239 | -3.8751 |
| 0.982186 | 0.736773 |
| 2.38993 | 0.392008 |
| 4.29965 | 3.06402 |
| -3.60466 | -6.80688 |
| 1.06331 | -1.38291 |
| 3.86222 | 0.852889 |
| 5.32853 | 5.95125 |
| -4.69174 | -8.62882 |
| -0.383243 | 1.53061 |
| 2.2503 | 0.717296 |
| 4.2965 | 7.65034 |
| -9.77993 | -7.43195 |
| -1.57704 | -1.68578 |
| 2.059 | 2.09355 |
| 4.72658 | 7.10818 |
| -3.65344 | -2.21279 |
| -0.544595 | 0.447653 |
| 2.47207 | 2.37372 |
| 4.28876 | 7.16518 |
| -7.75424 | -2.46897 |
| -0.922869 | -0.0177502 |
| 1.73353 | 1.01235 |
| 5.39667 | 5.76931 |
| -2.5157 | -6.15092 |
| 1.50383 | -1.29935 |
| 3.24336 | 0.475802 |
| 5.14904 | 5.97729 |
| -2.26732 | -3.33424 |
| 0.711436 | -0.736055 |
| 2.46248 | 0.540866 |
| 3.84866 | 4.63098 |
| -4.49551 | -7.32824 |
| 0.182898 | -1.76522 |
| 2.01987 | 1.14621 |
| 3.60407 | 7.43554 |
| -4.71859 | -2.54048 |
| -0.276082 | -0.700857 |
| 2.36996 | 0.891976 |
| 4.37185 | 7.95603 |
| -8.25623 | -0.118966 |
| -1.1639 | 3.1876 |
| 0.511214 | 1.31286 |
| 4.41716 | 5.71988 |
| -1.60728 | -1.86579 |
| 2.10699 | 0.933973 |
| 3.33054 | 3.0338 |
| 4.93409 | 6.16498 |
| -3.07983 | -6.73043 |
| 0.520754 | 1.1277 |
| 2.49201 | 3.1279 |
| 4.15259 | 8.44797 |
| -2.08505 | -6.36602 |
| 0.782842 | 0.79839 |
| 1.95131 | 2.53049 |
| 3.60355 | 5.27449 |
| -3.77761 | -4.49974 |
| 0.565573 | -0.256268 |
| 1.69447 | 1.58977 |
| 3.74154 | 4.82576 |
| -2.51115 | -5.63591 |
| -0.891623 | 1.27299 |
| 1.96086 | 1.74369 |
| 4.06452 | 4.44025 |
| -2.49841 | -2.71713 |

**Figure S3**. Observed versus predicted values of the daily NEE model.

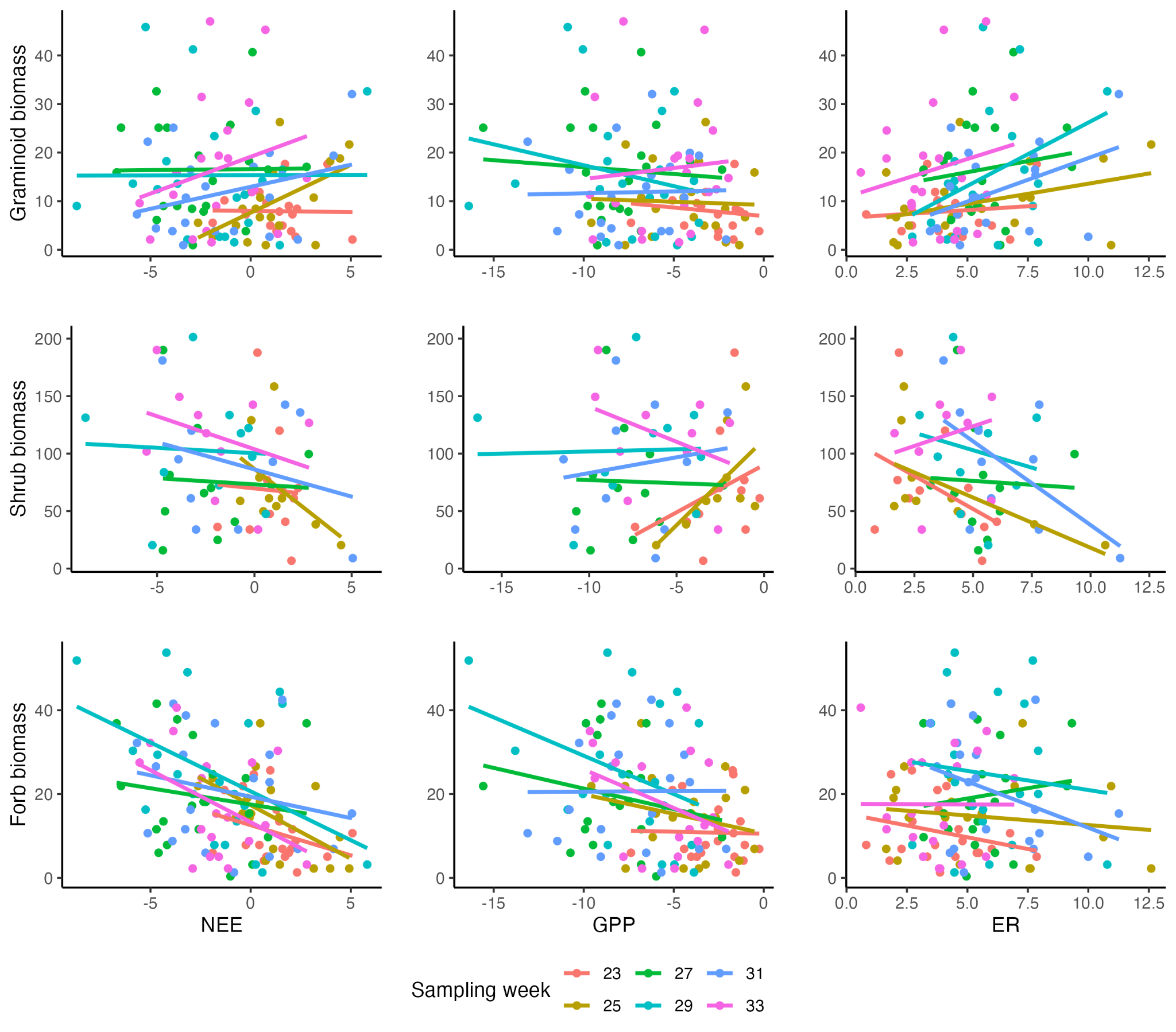

**Figure S4.** The linear relationships between carbon fluxes and graminoid, shrub, and forb biomass throughout the growing season. The dots and linear regression lines are color-coded by the week the data were collected.

**Tables**

**Table S1.** Species recorded and functional group classification.

| **Species name** | **Functional group** |
| --- | --- |
| *Dasiphora fructicosa* | Shrub |
| *Ribes montigenum* | Shrub |
| *Poa spp.* | Graminoid |
| *Festuca thurberi* | Graminoid |
| *Carex spp.* | Graminoid |
| *Achillea millefolium* | Forb |
| *Campanula rotundifolia* | Forb |
| *Collomia linearis* | Forb |
| *Delphinium nuttallanum* | Forb |
| *Erigerion speciosus* | Forb |
| *Erigeron elatior* | Forb |
| *Frasera speciosa* | Forb |
| *Galium septentrionale* | Forb |
| Helianthella quinquenervis | Forb |
| *Heuchera parvifolia* | Forb |
| *Hymenoxys hoopesii* | Forb |
| *Lathryus leucanthus* | Forb |
| *Ligusticum porteri* | Forb |
| *Linum lewisii* | Forb |
| *Potentilla pulcherrima* | Forb |
| *Senecio crassulus* | Forb |
| *Senecio spp.* | Forb |
| *Thalictrum fendleri* | Forb |
| *Tragopogon dubius* | Forb |
| *Vicia americana* | Forb |
| *Viola praemorsa* | Forb |

**Table S2.** The Bayesian model quantile credible intervals for all parameters. Tau is variance. In the treatment column, Treatment is the earlier snowmelt plots.

| **Parameter** | **Treatment** | **Season** | **2.5%** | **25%** | **50%** | **75%** | **97.5%** |
| --- | --- | --- | --- | --- | --- | --- | --- |
| Soil moisture | Control | Early | -0.063 | 0.066 | 0.146 | 0.229 | 0.370 |
| Soil moisture | Treatment | Early | -0.506 | -0.281 | -0.176 | -0.073 | 0.122 |
| Soil moisture | Control | Mid | -0.136 | -0.056 | -0.014 | 0.027 | 0.107 |
| Soil moisture | Treatment | Mid | -0.328 | -0.190 | -0.119 | -0.052 | 0.077 |
| Soil moisture | Control | Late | -0.341 | -0.207 | -0.138 | -0.068 | 0.058 |
| Soil moisture | Treatment | Late | -0.799 | -0.492 | -0.316 | -0.131 | 0.189 |
| Air temperature | Control | Early | -1.334 | -0.788 | -0.502 | -0.213 | 0.323 |
| Air temperature | Treatment | Early | -0.524 | 0.270 | 0.623 | 1.009 | 1.821 |
| Air temperature | Control | Mid | -0.178 | 0.031 | 0.157 | 0.280 | 0.527 |
| Air temperature | Treatment | Mid | -0.300 | 0.025 | 0.183 | 0.333 | 0.613 |
| Air temperature | Control | Late | -0.112 | 0.138 | 0.281 | 0.423 | 0.689 |
| Air temperature | Treatment | Late | -0.174 | 0.113 | 0.256 | 0.394 | 0.686 |
| Light | Control | Early | -0.003 | -0.002 | -0.002 | -0.001 | -0.001 |
| Light | Treatment | Early | -0.006 | -0.005 | -0.005 | -0.004 | -0.004 |
| Light | Control | Mid | -0.008 | -0.007 | -0.007 | -0.006 | -0.005 |
| Light | Treatment | Mid | -0.010 | -0.009 | -0.008 | -0.008 | -0.007 |
| Light | Control | Late | -0.008 | -0.007 | -0.007 | -0.006 | -0.005 |
| Light | Treatment | Late | -0.007 | -0.006 | -0.006 | -0.005 | -0.004 |
| Intercept | Control | NA | -2.869 | 0.587 | 2.351 | 4.168 | 7.159 |
| Intercept | Treatment | NA | -2.652 | 1.966 | 4.533 | 7.070 | 12.593 |
| tau | NA | NA | 0.161 | 0.183 | 0.195 | 0.208 | 0.234 |

**Table S3.** Linear mixed effect model results for the relationship between the different plant functional groups and carbon fluxes, with a bolded p-value indicating a significant effect.

| **Parameter** | **NEE** | | |  | **GPP** | | |  | **ER** | | |
| --- | --- | --- | --- | --- | --- | --- | --- | --- | --- | --- | --- |
|  | *df* | *F* | *p* |  | *df* | *F* | *p* |  | *df* | *F* | *p* |
| Treatment | 1,9 | 0.273 | 0.614 |  | 1,9 | 1.770 | 0.216 |  | 1,9 | 0.705 | 0.423 |
| Graminoid biomass | 1,38 | 0.251 | 0.619 |  | 1,38 | 5.434 | **0.025** |  | 1,38 | 2.123 | 0.153 |
| Shrub biomass | 1,38 | 9.947 | **0.003** |  | 1,38 | 4.632 | **0.038** |  | 1,38 | 1.173 | 0.286 |
| Forb biomass | 1,38 | 4.102 | **0.050** |  | 1,38 | 13.188 | **0.001** |  | 1,38 | 8.733 | **0.005** |
| Time | 1,38 | 4.145 | **0.049** |  | 1,38 | 9.797 | **0.003** |  | 1,38 | 6.177 | **0.018** |
| Treatment × Graminoid Biomass | 1,38 | 0.923 | 0.343 |  | 1,38 | 0.241 | 0.627 |  | 1,38 | 2.922 | 0.096 |
| Treatment × Shrub Biomass | 1,38 | 2.892 | 0.097 |  | 1,38 | 0.001 | 0.969 |  | 1,38 | 2.374 | 0.132 |
| Treatment × Forb Biomass | 1,38 | 0.619 | 0.436 |  | 1,38 | 0.566 | 0.456 |  | 1,38 | 0.291 | 0.593 |
| Time × Graminoid Biomass | 1,38 | 3.707 | 0.062 |  | 1,38 | 5.542 | **0.024** |  | 1,38 | 0.855 | 0.361 |
| Time × Shrub Biomass | 1,38 | 0.010 | 0.923 |  | 1,38 | 0.072 | 0.790 |  | 1,38 | 0.124 | 0.727 |
| Time × Forb Biomass | 1,38 | 0.001 | 0.973 |  | 1,38 | 0.068 | 0.796 |  | 1,38 | 0.124 | 0.727 |
